# Naturally occurring viruses of *Drosophila* reduce offspring number and lifespan

**DOI:** 10.1101/2023.09.05.555738

**Authors:** Megan A. Wallace, Darren J. Obbard

## Abstract

*Drosophila* remains a pre-eminent insect model system for host-virus interaction, but the host range and fitness consequences of the drosophilid virome are poorly understood. Metagenomic studies have reported over 160 viruses associated with *Drosophilidae*, but few isolates are available to characterise the *Drosophila* immune response, and most characterisation has relied on injection and systemic infection. Here we use a more natural infection route to characterise the fitness effects of infection and to study a wider range of viruses. We exposed laboratory *D. melanogaster* to 23 naturally occurring viruses from wild-collected drosophilids. We recorded transmission rates along with two components of female fitness: survival and the lifetime number of adult offspring produced. Nine different viruses transmitted during contact with laboratory *D. melanogaster*, although for the majority, rates of transmission were less than 20%. Five virus infections led to a significant decrease in lifespan (D. melanogaster nora virus, D. immigrans nora virus, Muthill virus, galbut virus and Prestney Burn virus), and three led to a reduction in the total number of offspring. Our findings demonstrate the utility of the *Drosophila* model for community-level studies of host-virus interactions, and suggest that viral infection could be a substantial fitness burden on wild flies.

## 1 Introduction

Insects and their viruses are often assumed to be involved in reciprocal, co-evolutionary arms-race dynamics [1]. This is broadly supported by the fast evolution of some insect immune genes [2], the presence of large-effect polymorphisms for resistance [3,4] and the evolution of viral suppressors of immunity [5]. However, the strength of virus-driven selection on immune genes depends on the fitness effects of infection, and the costs imposed by the majority of insect viruses are unknown. One way to assess the importance of virus-mediated selection in insects is to characterise viral fitness costs in a well-studied model system such as *Drosophila*.

Over 160 *Drosophila*-associated viruses have now been reported across the genus [6–10]. Metagenomic sequencing has also revealed overlapping host ranges for many *Drosophila* viruses [6–9,11]. This suggests that the cross-species transmission may be common in the wild, and creates an opportunity to study virus-driven selection in a multi-host, multi-virus community. However, less than 10% of these viruses have been isolated—usually the first step to characterising fitness effects. Thus, our dependence on the few isolated, well-characterised viruses for experimental studies may give us a biased view of virus-mediated selection.

The nine isolated *Drosophila* viruses include seven RNA viruses: D. melanogaster sigmavirus (DmelSV), Drosophila C virus (DCV), Drosophila A virus (DAV), D. melanogaster nora virus (DmelNV), La Jolla virus, Newfield virus and galbut virus [12–18] and two DNA viruses: Drosophila innubila Nudivirus (DiNV) and Drosophila Kallithea Nudivirus (KV) [6,19]. Infection with some of these viruses has large fitness costs. For example, injection with DCV (+ssRNA, *Dicistroviridae*) leads to a systemic infection that causes intestinal obstruction, impaired locomotor activity, metabolic depression, and eventual mortality in *D. melanogaster* adults [20,21]. Systemic infection with DiNV (dsDNA, *Nudiviridae*) causes a decrease in lifespan in both *D. innubila* and *D. falleni*, and infection in wild females of both species incurs an ∼80% reduction in offspring production [6]. Systemic infection with KV, another large dsDNA Nudivirus, into *D. melanogaster* causes increased male mortality, and for females, decreased movement and late-life egg laying [19]. Recently, reductions in egg laying were also reported in four *Drosophila* species on infection with two RNA viruses: Newfield virus (41-94% reduction) and La Jolla virus (8-91% reduction) [18].

In contrast to these clear disease phenotypes, infection with the most common virus in wild Dmel populations – galbut virus – seems in previous studies to cause no significant decrease in average lifespan or total offspring production [22]. Other isolated *Drosophila* viruses have subtle pathologies, but can still result in a sizeable reduction in fitness. This makes characterising their fitness costs challenging even when a viral isolate is readily available. For example, the vertically transmitted DmelSV causes no detectable decrease in adult mortality, but does reduce fitness by an estimated ∼25% [23,24]. Persistent infection with DmelNV (+ssRNA, *unclassified Picorna-like*) also produces subtle pathologies: an increased vacuolization of the intestine [25], and decreased climbing ability [26]. Indeed, until recently it was believed that, before 50 days, the survival of DmelNV infected flies was similar to that of uninfected flies [25]. However, in contrast to these results, a recent study found that DmelNV presence in some lab stocks reduced lifespan by >40% [27]. For DAV (+ssRNA), which is widespread and replicates in laboratory flies and cell culture [28,29], inoculation with the virus only seems to increase mortality >30 days post infection, but the virus can still reduce fecundity by 28% on oral infection [4]. However, DAV clearly reduces lifespan in *Drosophila suzukii* [16], consistent with the idea that virulence varies among *Drosophila* host species [30,31].

Characterisation of the natural fitness costs of *Drosophila*-virus infection is also hampered by a reliance on injection into the thorax or abdomen—transmission routes that are thought to be rare in the wild [see methods in 32]. These methods facilitate laboratory studies and standardise doses, but may not replicate natural fitness costs as infection phenotypes often vary depending on infection route. Compared to systemic injection, oral infection—which is likely to be more common in the wild—can infect different tissues [33], produce different disease phenotypes [34,35] and induce different immune responses [36, and see review 37].

Directly comparing the fitness of naturally-infected and uninfected wild individuals avoids problems associated with systemic injection, allows use of non-isolated viruses, and provides the most accurate picture of the strength of virus-driven selection in natural populations. Nevertheless, few papers have used this approach [but see 6] as it is difficult to ensure sufficient statistical power to detect effects. To remedy this, forced contact in the lab can create opportunity for viral transmission between infected and uninfected individuals, allowing the virus to transmit using its natural route while avoiding the need for a viral isolate or systemic injection.

In this study, we use contact between infected and uninfected individuals to infect *D. melanogaster* females with both native viruses and viruses carried by other species of *Drosophila*. We assessed the effect of viral infection on two key components of fitness: lifespan and offspring production. We find that many *Drosophila* viruses transmit readily across species, in line with previous observations of host range overlap. Additionally, we identify five viruses associated with a significant reduction in lifespan, and three with a reduction in offspring production in *D.melanogaster* females. These reductions to key components of individual fitness could create significant selection pressure on the hosts, and potentially drive the evolution of immune genes.

## 2 Methods

### 2.1 Exposure of D. melanogaster to wild Drosophila

To detect fitness costs associated with viral infection, we exposed laboratory *D. melanogaster* to viruses carried by wild-collected flies, and followed the subsequent lifespan and offspring production of the laboratory flies. Wild ‘donor’ (source) flies were collected between the 2^nd^ and 8^th^ July 2017 from banana/yeast baits in Sussex (England; 51.100N, 0.164E), and Edinburgh (Scotland; 55.919N, -3.211W). We collected male and female *D. melanogaster*, *Drosophila immigrans*, and three species of the *Drosophila obscura* species group (*Drosophila obscura*, *subobscura*, and *subsilvestris*). However, members of the obscura group were treated together, as females are difficult to distinguish morphologically and carry partly overlapping viral communities [8]. The wild flies, separated by species, were then kept at high density (8-20 flies per vial) on solid sugar/agar medium for 2-7 days.

We prepared laboratory ‘recipient’ flies from *Wolbachia*-negative laboratory ‘Oregon R’ *D. melanogaster* (see supplementary methods). Briefly, we collected flies one day after eclosion and 43 groups of 25 females and 10 males were permitted to mate on sugar/agar vials for four days. Twenty females from each vial were then moved to one of the vials containing wild donor flies of one of three different species (obscura group, *D. immigrans*, and *D. melanogaster*), so that these co-housing vials contained 10 wild flies of one species and 20 laboratory *D. melanogaster* females. In the case of *D. melanogaster*, only male wild donors were used, to enable wild and laboratory flies to be subsequently distinguished. To partly control for other impacts of the exposure period, in a fourth treatment we co-housed laboratory females at the same density with males of the same laboratory stock that were not expected to carry any of the viruses seen in the wild flies. After a three-day exposure period to permit virus transmission, we removed the wild donors and laboratory males (in the control vials) and froze them at -80C for virus RT-PCR assays. At this point we were necessarily blind to the viruses present (if any) and their transmission. We then placed the female laboratory recipient flies individually into vials containing Lewis *Drosophila* medium (see supplementary methods) along with two fresh laboratory male flies to ensure continued mating opportunity.

We observed the ‘recipient’ laboratory females on a daily basis for the remainder of their lives, tipping them on to fresh medium on days 2, 5, 10, and then at 7 day intervals thereafter for a maximum of 72 days. Dead females were removed and frozen at -80°C for later virus assays, and their day of death recorded. As a proxy for fecundity, we recorded the number of eclosed adult offspring laid by each female during the 15 days after exposure. Throughout the experiment, we immediately replaced dead males with fresh individuals to maintain the same rate of courtship and mating. Note that the only difference between the routine maintenance of laboratory flies and the treatment of the recipient laboratory flies was the three-day period of exposure to wild *Drosophila* in their first week as adults.

### 2.2 Detection of virus infection and transmission

We extracted RNA from the wild donor flies and laboratory recipient flies using Tri-Reagent Solution (ThermoFisher Scientific) according to the manufactures protocol. We synthesised cDNA with M-MVL reverse transcriptase (Promega) and random hexamer primers (ThermoFisher Scientific; see supplementary methods). We then used RT-PCR assays to reveal the viruses to which laboratory recipient females had been exposed, testing each of the wild-collected donor groups (*D. immigrans*, *D. melanogaster*, obscura group) and the control laboratory *D. melanogaster* males for a total of 59 different viruses (see table S1 for the primers; and supplementary methods for further detail). Four of the detected viruses (Drosophila Crammond virga-like virus, Drosophila Sighthill phlebovirus, Drosophila sunshine bunyavirus and Drosophila Burdiehouse burn chuvirus) were novel viruses recently described from metagenomic sequencing of the obscura group, *D. immigrans*, *D. melanogaster* and *D. funebris* [38].

We determined which viruses had transmitted to the laboratory *D. melanogaster* by assaying the dead recipient females for those viruses they had been exposed to, based on the viruses detected in the donor vial. Because we harvested the recipient flies on the day they died naturally, usually many days after the last exposure to the donor flies (1-72 days, mean 34), it is unlikely that inactive viral material remained as surface or gut contamination. However, a delay of up to 24 hours between death and preservation at -80°C may also have led to substantial RNA degradation—although this is likely to be mitigated for encapsidated virions. Thus, to maximise the chance of virus detection we designed additional PCR assays with short amplicon lengths (120-170bp) using the Primer3 [39] primer design tool in Geneious v10.1.3 (http://www.geneious.com/; primers provided in table S2). Note that misclassification of infection status in the recipient flies, such as false negatives from virus detection, will tend underestimate of the impact of infection by erroneously reducing the distinction between infected and uninfected recipients. For galbut virus we used multiple primers (for segment 1 and for the optional or satellite ’chaq’ segment; Genbank accession: KP714088). There were four occasions on which the chaq PCR gave a positive result but the segment 1 PCR did not. We considered these to be PCR dropouts, and treated these flies as positive for galbut and chaq.

For each wild collected donor species, we recorded the viruses present in each of the co-housing vials and those transmitted to recipient *D. melanogaster* females. As we assayed each vial of donor flies as a pool, we could not estimate individual-level prevalence directly for the donor species. We therefore estimated global virus prevalence in these flies by maximum-likelihood, assuming pooled Bernoulli trials [40]. We quantified transmission rate for each specific host:virus combination by calculating the number of recipient individuals infected with a virus as a proportion of the number exposed. Three vials of laboratory *D. melanogaster* males unexpectedly tested positive for virus infection (one for Muthill virus and two for DimmNV), perhaps suggesting a low level of viruses circulating in the laboratory. However, as the primary comparison in our study is between the recipient flies that were infected and those that were not, we were able to treat these exposures in the same way as exposure to viruses from wild flies.

### 2.3 Sequence confirmation of transmitted viruses

For some viruses, such as those thought to be exclusively vertical in their transmission or limited to a single host species, we chose to perform additional confirmation of virus identification and classification, and thus rule out the possibility of cross priming with another related virus. We did this in two ways. First, by Sanger sequencing short (600nt) RT-PCR products of the RNA-dependent RNA polymerase (RdRp) and Viral protein 2 (VP2) of DimmNV, and the L (∼630bp) and N (∼525bp) genes of DimmSV. We sequenced these regions from a sample of both the wild collected donor flies and the infected recipient laboratory *D. melanogaster* females (see supplementary methods). Second, we selected and pooled RNA from six infected recipient flies for meta-transcriptomic sequencing, three infected with DimmSV, and three with DimmNV. We extracted total RNA as previously described for all other recipients. A single sequencing library for the pool was prepared from the NEBNext Ultra™ RNA Library Prep Kit for Illumina (New England Biosciences), with no poly A selection, and ribosomal RNA depleted using primers specifically designed against *Drosophila* rRNA. The pooled sample was then sequenced at the University of Liverpool centre for genomics research on an Illumina NovaSeq 6000 using S4 chemistry, producing 30 million 150 nt read pairs.

### 2.4 Analysis of lifespan and offspring production

We analysed survival and fecundity data using Bayesian Generalized Linear Mixed Models (GLMMs), implemented in the package MCMCglmm [41] in R v.4.0.3 [42]. For our longevity analysis, we fitted two alternative parameterisations. First, we fitted the presence/absence of each virus as a fixed effect (9 factors with two levels), with exposure vial as a random effect to model any other sources of among-vial variance in mortality:

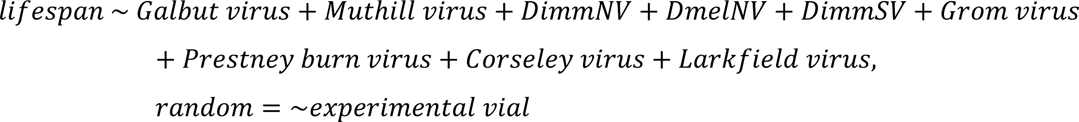

Second, we alternatively modelled the number of viral infections as a continuous predictor with virus identity as a random effect. This second model allowed us to ask whether the virus identity caused a significant deviation in lifespan from the general ‘virus effect’, and whether viral infections have an additive effect on lifespan when co-infecting individuals (syntax as for MCMCglmm):

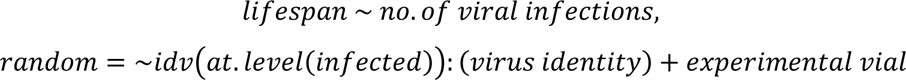

In this model, idv() denotes the form of the variance structure, fitting a single variance that is shared by all viruses in infected individuals. For both of these analyses, we modelled random effects and residual variances using Gaussian distributions and we confirmed that the model residuals appeared approximately Gaussian. For each analysis, we ran the Markov chain for 1×10^6^ steps, and recorded 10,000 of these, ensuring all parameter estimates had effective sample sizes of > 5,000.

We modelled the lifetime number of offspring using a hurdle-Poisson distribution to account for the substantial number of females who produced no offspring. This approach models two latent variables: (1) the number of offspring from each individual (if any), which is assumed to follow a zero-truncated Poisson distribution, and (2) the probability of zero offspring from an individual (on the logit scale). As for the first lifespan analysis above, we modelled the presence/absence of each virus as a fixed effect, and the exposure vial as a random effect. Note that for two of the viruses all infected females produced offspring, such that the probability of zero offspring for those viruses could not be estimated. The variance structure specified using idh() permits the Poisson mean and the probability of zero offspring to differ in their variance, and sets the covariance between them to zero. We supressed the overall intercept of the model, so that we could interpret the estimates for the probability of zero offspring production independently of the Poisson model estimates:

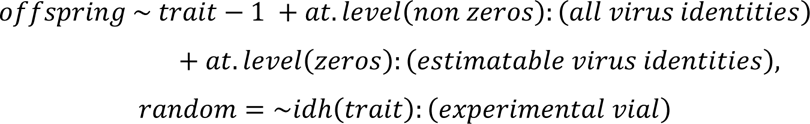

We then simulated data based on our posterior estimates to confirm that our parameter estimates were consistent with the observed number of zeros in the data (Fig. S1). We recorded 20,000 steps in the Markov Chain, ensuring all parameter estimates had effective sample sizes of >1,500 In addition, because a reduction in total offspring number might reflect either reduced lifespan (as analysed above) or a reduced rate of egg laying, we also separately analysed early offspring production (first five days post exposure, i.e. to 13 days post-eclosion). In this analysis, we excluded 10 flies that died prior to day 13, so that there is no direct impact of lifespan on offspring production. We fitted the model exactly as for lifetime offspring production (above), except that the probability of zero offspring could not be estimated for five of the viruses. The sample size of the chain was 200,000 and all posterior samples had an effective sample size of > 4,000.

## 3 Results

### 3.1 Host range, prevalence, and transmission

Using RT-PCR we identified 23 different viruses carried by 430 wild-collected *D. immigrans*, *D. melanogaster*, and obscura-group flies (Fig. 1, and Table S3). Two viruses were found in all three species groups: *D. immigrans* nora virus (DimmNV), which is a +ssRNA *picorna*-like virus related to members of *Orthonoravirus* in the *Noravirirdae*; and Muthill virus, a +ssRNA *Nege*-like virus. Each of the other viruses occurred in a single wild species group. Our estimates of wild virus prevalence ranged from a maximum of 100% (2 Log-likelihood bounds, 84-100%) for galbut virus in *D. melanogaster* to a minimum of 0.69% (0.7-3.8%), for Blackford virus, Corseley virus, Crammond virus, DimmNV, Muthill virus and Pow Burn virus in the obscura group—the latter consistent with a single infected fly in our collection. The high prevalence of galbut virus in our *D. melanogaster* sample is consistent with its previously reported high prevalence in the wild in [7], and in recently established colonies in [17].

**Fig. 1:**
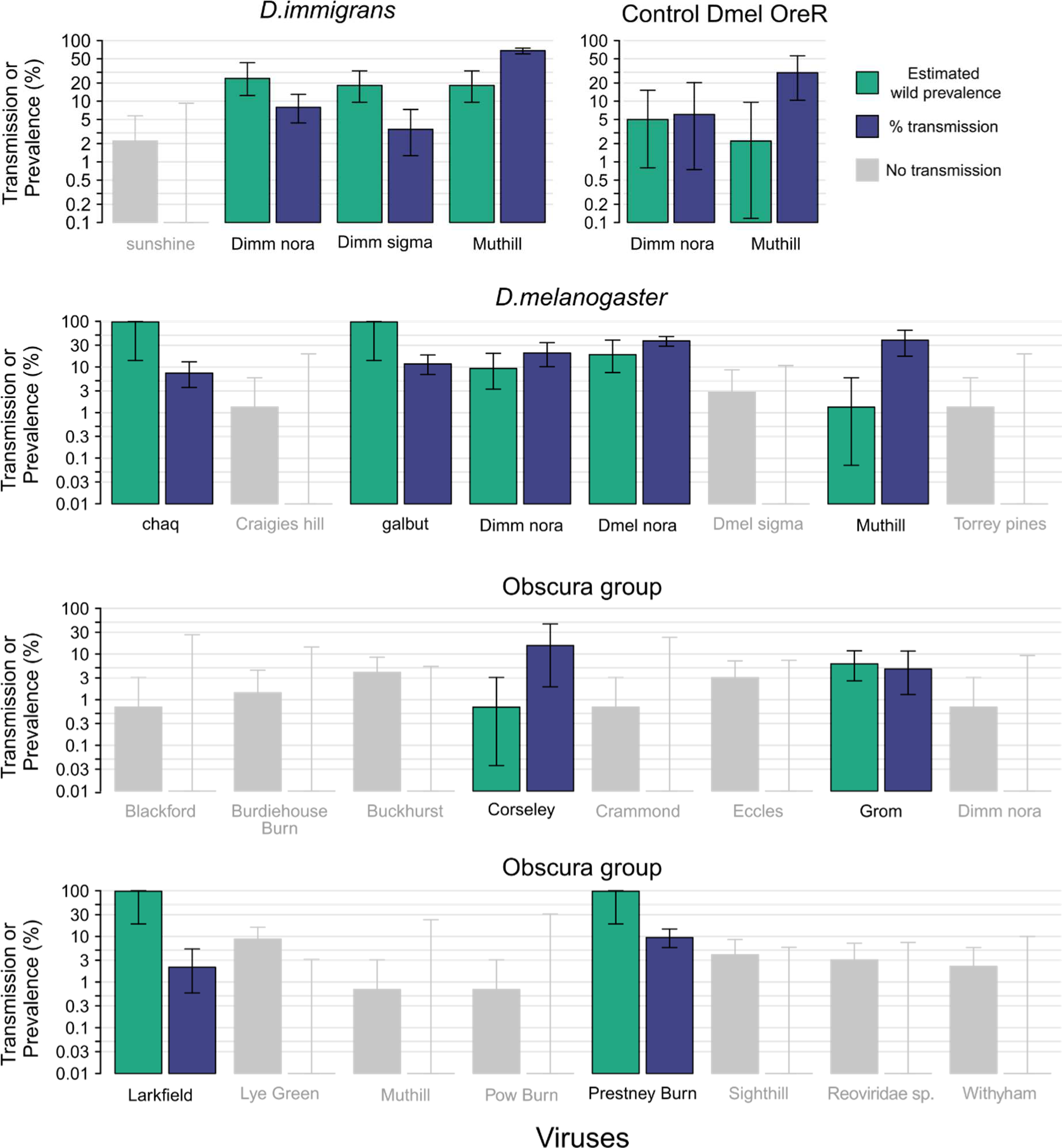
Summary of viral exposure and transmission. Maximum likelihood estimates of viral prevalence (green if transmitted, or grey if not transmitted) and transmission (purple) of viruses from wild-collected ‘donor’ flies to ‘recipient’ laboratory *D. melanogaster* during three days of exposure, assayed at time of death. Error bars show 2 log-likelihood intervals. Note the logarithmic y-axis, and that ‘chaq’ is thought to be either an optional segment or satellite virus of galbut virus [17].

Nine viruses were detectable in the exposed laboratory *D. melanogaster* at their time of death (galbut virus, DmelNV, DimmNV, DimmSV, Grom virus, Prestney burn virus, Larkfield virus and Corseley virus), suggesting that they had been infected during exposure to wild flies. There was no detectable difference among donor groups in the in the rate of transmission (3/4 viruses from *D. immigrans*, 4/7 from *D. melanogaster*, 4/16 from Dobs; *p*=0.116 by Fisher’s Exact Test, although power to detect this was low and it may be notable that few viruses were transmitted from the obscura group).

Some host species:virus combinations produced a consistently high rate of transmission. For example, over 35% of the *D. melanogaster* exposed to Muthill virus from *D. melanogaster* or *D. immigrans* were infected (68% and 39% from *D. immigrans* and *D. melanogaster* respectively). Moreover, these infections persisted for a substantial period, as flies testing positive for Muthill virus survived an average of 38.4 days. Nevertheless, the vast majority of viruses transmitted to <15% of the exposed individuals, leading to relatively low power to detect the effect of viral infection on fitness.

For some viruses, the observed natural infections of *D. melanogaster* were unexpected. For example, DimmNV has not previously been reported to infect *D. melanogaster* [7] (see also [43]). Whereas here we observed both natural infections in the field (estimate of wild viral prevalence, 9%, 2 Log-likelihood bounds, 3-20%) and transmission in the lab (transmission rate from wild *D. melanogaster*, 20.4%; from wild *D. immigrans*, 7.9%). Nevertheless, sequencing was able to confirm the identity of these infections, with putative DimmNV sequences from wild donor flies and a recipient *D. melanogaster* being >99.5% identical to VP2 (RdRp) of published DimmNV (KF242511.1 [43]), and with DimmNV from additional wild-collected flies from the same geographic location showing no detectable differentiation between *D. melanogaster* and *D. immigrans* (S1 & Fig S2).

Perhaps even more surprisingly, given the usually vertical transmission of sigma viruses [44–46], we also observed apparent transmission of DimmSV from wild *D. immigrans* into six recipient laboratory *D. melanogaster* females (i.e. 3.4% of the recipient flies that were exposed). To test for the possibility that these sequence reads may represent gut or surface contamination (albeit a mean of 19.8 days after a three day exposure as adults to *D. immigrans*), we conducted pooled RNA sequencing on the six *D. melanogaster* females that tested positive for DimmSV by RT-PCR. This showed that small amounts of *D. immigrans* RNA was present in these samples, suggesting that these apparent transmissions by DimmSV may represent cross contamination rather than infection. Note, however, that such contamination in general would reduce the apparent fitness impact of viral infection (by misclassifying uninfected flies as infected).

Galbut virus, a partitivirus that was recently found to have efficient bi-parental vertical transmission [17], also appeared to transmit from wild *D. melanogaster* males into 16 recipient laboratory *D. melanogaster* females (i.e. 11.8% of the recipient flies that were exposed). Its satellite—chaq—was transmitted to 10 of these 16 individuals, consistent with previous observations of its presence in most, but not all, individuals infected with galbut virus [17]. However, in a previous study, levels of galbut virus RNA in flies 21 days after virus ingestion were thought to be inconsistent with viral replication or horizontal transmission. Alternatively, it is possible that these reads derive from vertically transmitted infections in the embryos within the sequenced mother. Thus, despite the persistence of galbut virus RNA in flies long after exposure (mean of 23.4 days in our experiment), evidence of transmission remains equivocal.

### 3.2 Five naturally-occurring *Drosophila* viruses significantly reduce lifespan in *D. melanogaster*

We sought to detect an association between lifespan and virus infection status at the time of death. By comparing the 213 virus-infected *D. melanogaster* with the 393 uninfected individuals exposed at the same time in the same vials, we found a substantial reduction in lifespan associated with some viral infections (Fig. 2). In the fixed-effects model, five viruses were each significantly associated with reduced lifespan: galbut virus, DmelNV, DimmNV, Muthill virus and Prestney burn virus (Fig. 3). The viruses associated with the largest reduction in lifespan were DimmNV and Prestney burn virus, which appeared to reduce lifespan by 30% (95% Highest Posterior Density credibility interval: 15 to 46%) and 37% (95% CI 19 to 54%) respectively. DmelNV, Muthill, and galbut virus were associated with a reduction in lifespan of 17% (95% CI 5 to 28%), 11% (95% CI 4 to 18%), and 23% (95% CI 4 to 41%), respectively (Table S4). In this model there was no additional detectable variance in survival associated with exposure vial (95% CI 0 to 9% of the variance).

**Fig. 2:**
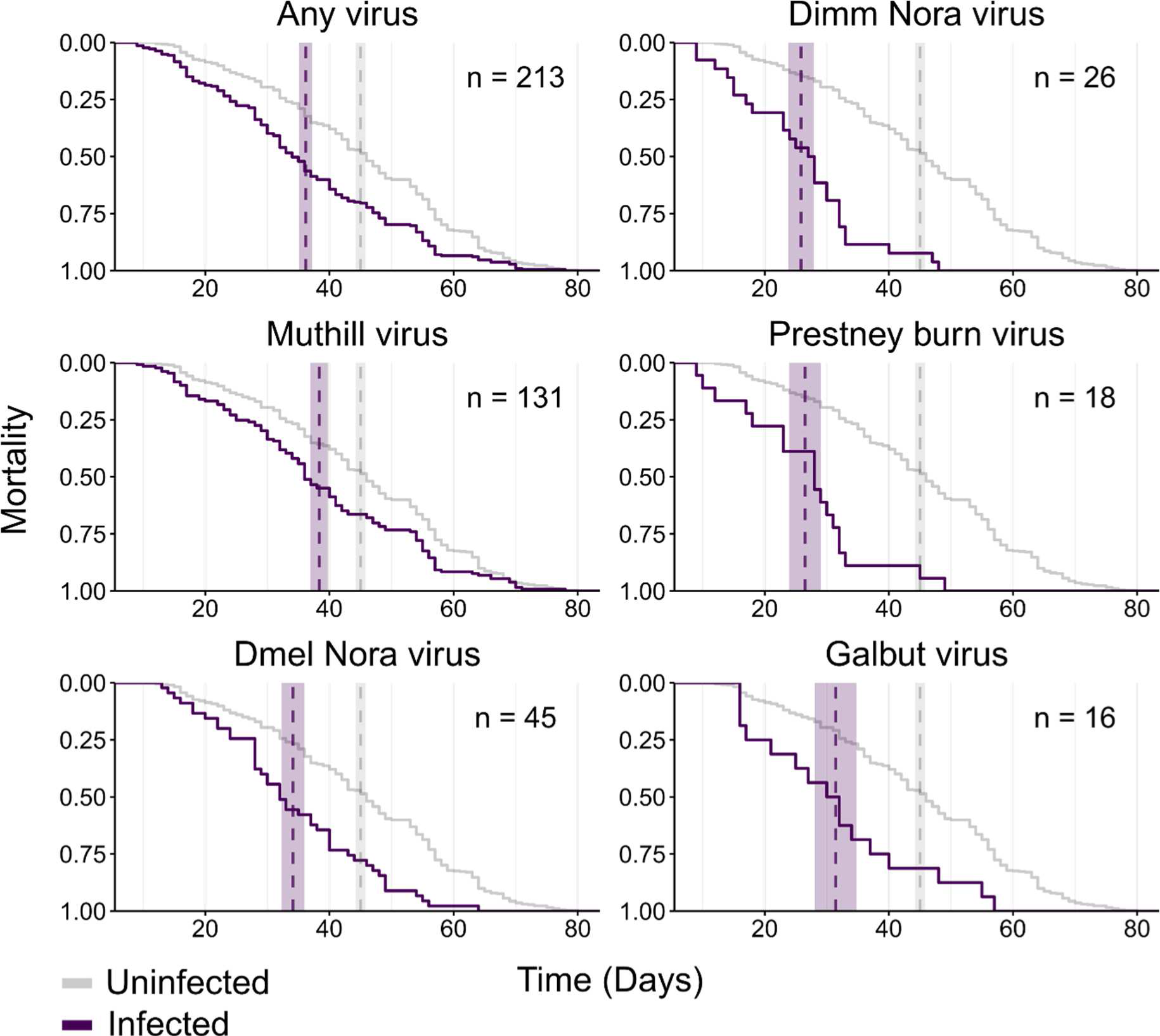
The lifespan of flies according to infection status. Panels show the mortality curves of infected (purple) and uninfected (grey line) laboratory *D. melanogaster* females. The number of infected flies is shown for each virus (n) and the dashed vertical lines indicate the mean lifespan (shaded areas give ±1 standard error). Note that the same 393 uninfected flies are depicted in each graph for comparison. Only those viruses with at least 10 infections are shown.

**Fig. 3:**
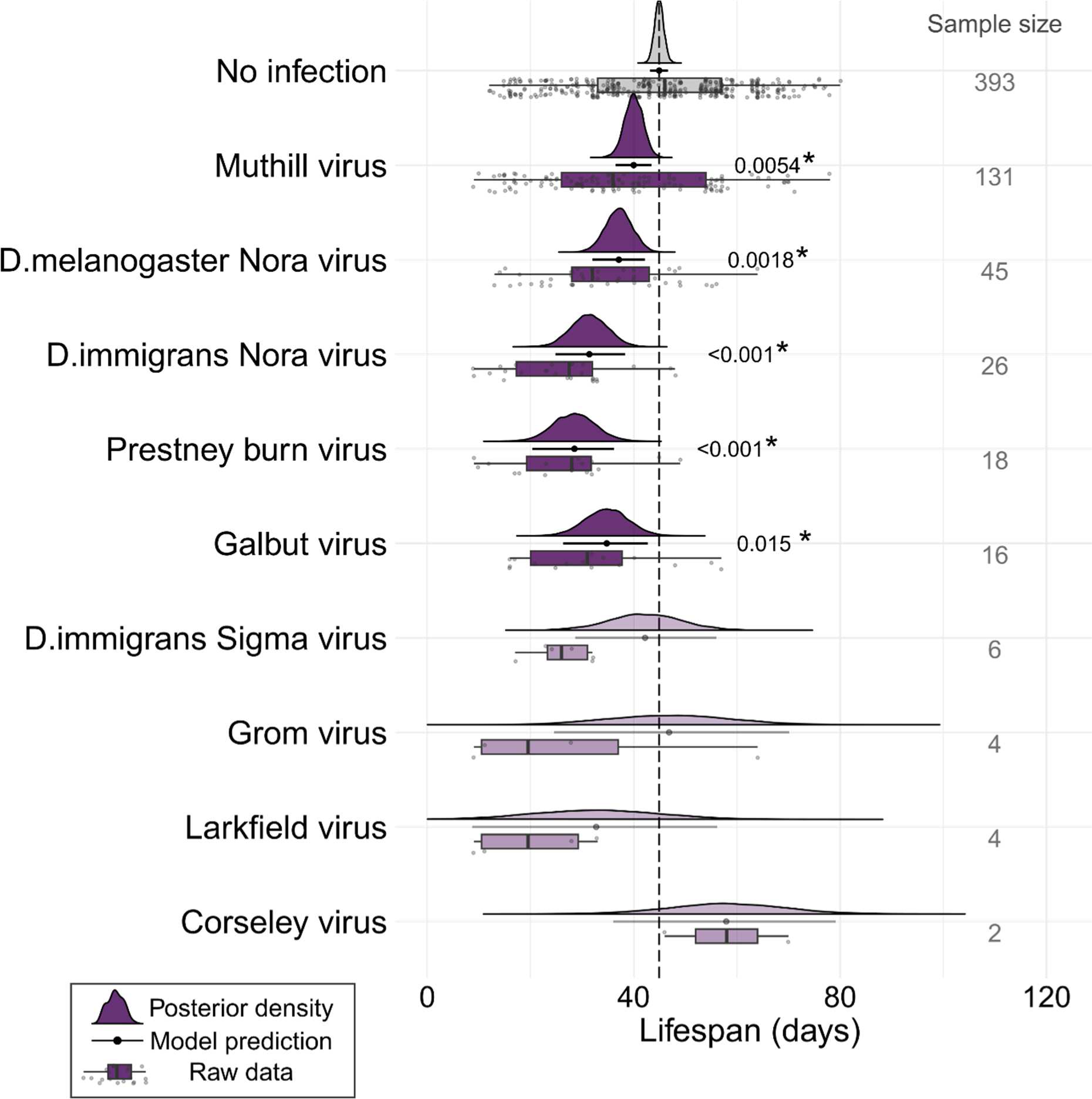
Posterior estimates of lifespan in infected flies. Plots show the output of the model in which virus identity was fitted as a fixed effect. Rows show the lifespan of *D. melanogaster* females that either did not test positive for infection (top row), or tested positive for each of nine different viruses (subsequent rows). Boxplots and points indicate the raw data with median and inter-quartile range. Filled curves show the posterior density of estimates, with solid horizontal lines and points to indicate the model prediction and 95% HPD credibility intervals. Sample size indicates the number of individuals that tested positive for each virus. For the alternative random-effects model, and a comparison between the model predictions see Fig. S4

Although the other viruses were not individually significantly associated with reduced lifespan, for all but two viruses the point estimate was a reduction rather than an increase in longevity (and power was low, with as few as two infections—in the case of Corseley virus). Thus, to test whether there was an overall association between lifespan and viral infection, and to test whether the effect of the number of infections was cumulative, we also fitted an alternative model in which we used the number of infections as a continuous predictor (zero being uninfected) and virus identity as a random effect (Fig. S4). This model showed that an increased number of viral infections was associated with a greater reduction in lifespan, with a reduction of 7.1 days for each additional virus detected in the fly (95% CI 2.7 to 10.8 days) (Table S4 & Fig. S5). However, specific virus identities were not associated with any detectable variance in lifespan (95% CI 0 to 24% of the variance). Taken together, these two alternative analyses suggest that there is a significant and substantial reduction in lifespan in those flies that test positive for a virus, but that while this is detectable for the most commonly transmitted viruses (and for all viruses considered together), it is not possible to separate the effect of less frequently transmitted viruses. For further comparison between these models, see supplementary materials.

### 3.3 Three naturally-occurring *Drosophila* viruses significantly reduce offspring number in *D. melanogaster*

Those *D. melanogaster* females that tested positive for a virus after death produced a mean of only 75 adult offspring during their lives, as compared to 102 offspring for those that did not test positive (Fig. S6). We analysed these data using a hurdle-Poisson GLMM to ask whether specific viral infections were associated with a decrease in the probability of having offspring, or in the mean number of offspring produced. Infection with three viruses, DmelNV, DimmNV and Muthill virus, was significantly associated with a reduced number of offspring produced, conditional on there being some offspring. Most notably, DmelNV infection was associated with a decrease in lifetime offspring production of 46% (95% CI 29 to 63%). DimmNV and Muthill virus were associated with a decrease in lifetime offspring production of 39% (CI 8 to 65%) and 24% (CI 9 to 39%) respectively. For all viruses, the point estimates were consistent with a reduction, rather than increase, in offspring number. In addition, Muthill virus was significantly associated with an increased likelihood of producing zero offspring, and again the (non-significant) point estimates for the effect of infection on sterility were similar for many other viruses. See Figure 4 & Table S5 for a summary. As for lifespan above, exposure vial did not explain a significant proportion of the variance in reproductive success (95% CI: 0 to 5% of the variance).

**Fig. 4:**
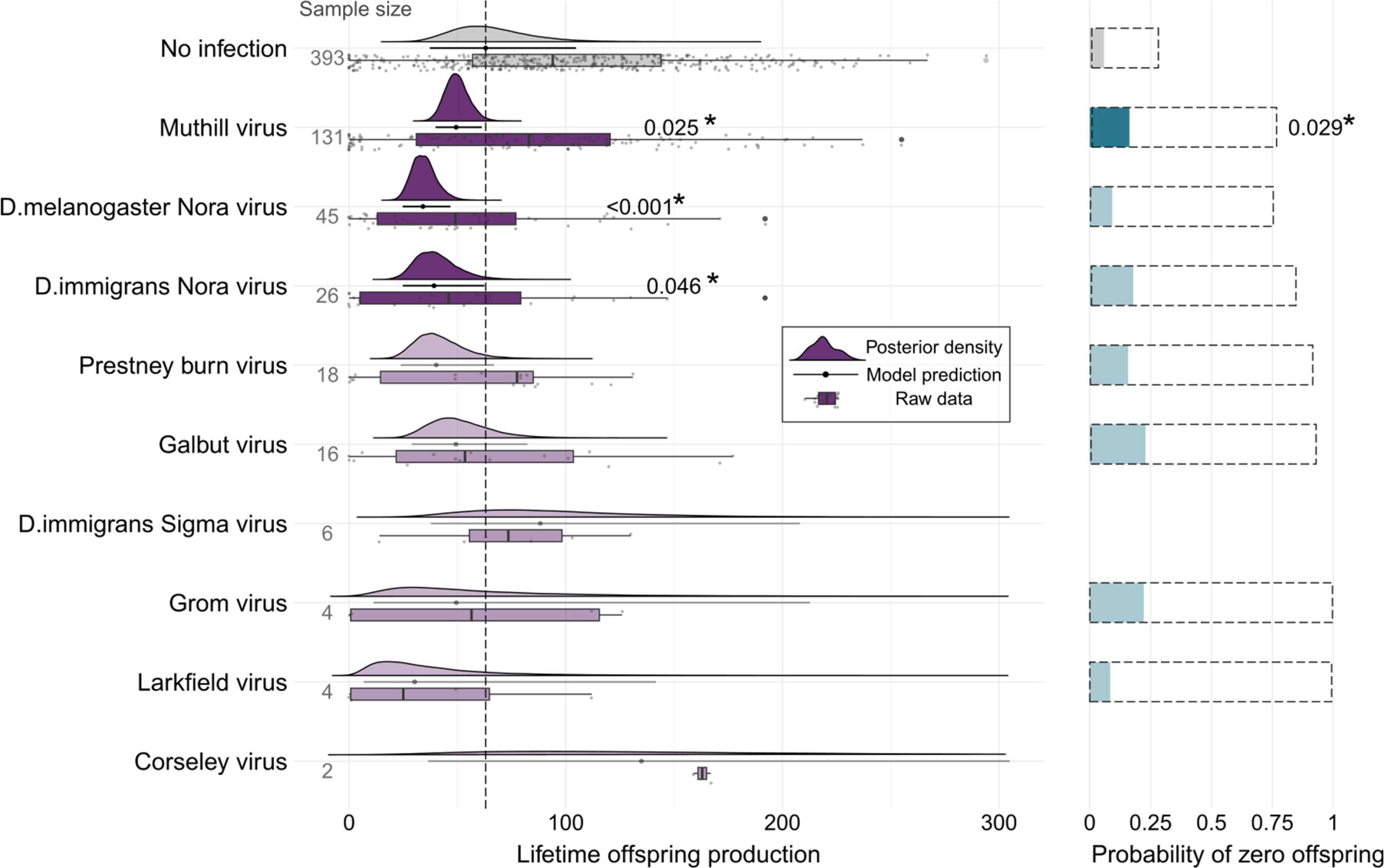
The impact of viral infection on lifetime offspring production in female *D. melanogaster* (OreR). Rows show the offspring production of *D. melanogaster* females that either did not test positive for infection (top row), or tested positive for each of nine different viruses (subsequent rows). Boxplots and points indicate the raw data with median and inter-quartile range. Filled curves on the left show the posterior density of estimates for the non-zero Poisson number of offspring, with solid horizontal lines and points to indicate the model prediction and 95% HPD credibility intervals. Sample size indicates the number of individuals that tested positive for each virus. Horizontal bar plots on the right show the predicted probability of zero offspring being produced with dashed line boxes to indicate the 95% HPD credibility intervals on these estimates.

A reduction in total offspring number might simply be a direct consequence of reduced lifespan, or it might also reflect a lower rate of offspring production. Therefore, we also examined the association between viral infection and the number of adult offspring produced during the first 5 days of egg-laying (females up to 13 days old). This was prior to any substantial mortality, and the analysis excluded the 10 females that died before this time point. Offspring production before day 5 of the experiment was significantly correlated with lifetime offspring production (Pearson’s product moment correlation = 0.55; 95% confidence interval = 0.49–0.60, t = 16.11, p = < 2.2e-16). We again found that DmelNV was significantly associated with a reduction in offspring production (Poisson regression coefficient -0.37, 95% HPD CI -0.7 to -0.05; pMCMC = 0.025*), showing that, for this virus, the reduction in offspring number is unlikely to be wholly attributable to reduced lifespan. In addition, galbut virus was associated with an increased likelihood of producing zero offspring in early-life, which seems to be compensated for when whole lifetime offspring production is analysed (Fig. S7 & Table S6).

## 4 Discussion

### 4.1 Cross-species transmission of *Drosophila* viruses

Many experiments that characterise *Drosophila* virus pathology use systemic injection. This limits experiments to isolated viruses, and may affect infection outcomes differently than natural transmission routes [see review 37]. Here we bypassed virus isolation or injection, and instead simply exposed laboratory *D. melanogaster* females to wild flies, later detecting transmission by RT-PCR. In total, our wild-collected ‘donor’ flies carried 23 viruses, with virus prevalence and host range broadly consistent with earlier surveys [8]. Nine viruses were transmitted to *D. melanogaster* during a three-day exposure window and remained detectable at the fly’s death. Transmission occurred even between the distantly-related hosts *D. melanogaster* and *D. immigrans*, which share a common ancestor ∼45 million years ago [47]. This suggests that many *Drosophila* viruses have a broad host range and can transmit among species by simple contact.

DimmNV (+ssRNA, *unclassified Picorna-like,* related to *Nodaviridae*) and Muthill virus (+ssRNA *Nege*-like virus) showed a particular propensity to infect multiple species and transmit to *D. melanogaster*. DimmNV is a close relative of DmelNV, which infects the midgut of laboratory and wild *D. melanogaster* [33], and has faecal-oral transmission [25]. However, DimmNV encodes a viral suppressor of RNAi (VSR) that—unlike the one encoded by DmelNV—is unable to supress the activity of *D. melanogaster* Argonaute-2 [43]. Despite this, here we found a 9% prevalence of DimmNV in wild *D. melanogaster* (2 log likelihood bounds = 6 - 29%), and a high rate of transmission from *D. immigrans* to *D. melanogaster* (18% of exposed individuals). Additionally, we recently described a ∼30% DimmNV prevalence in local *D. melanogaster* populations [38], and have identified small numbers of DimmNV reads in pools of wild-collected *D. melanogaster* [48] and *D. suzukii* [9] which have no contaminating *D. immigrans* reads (290 DimmNV reads in *D.suzukii,* 54 in *D.melanogaster* – Fig. S14). Together, these observations suggest that efficient activity of this VSR is not essential for natural DimmNV infection.

### 4.2 Possible horizontal transmission of putatively vertically transmitted viruses

We also unexpectedly observed horizontal transmission of two putatively vertically transmitted viruses: DimmSV to 3.4% of exposed females, and galbut virus to 11.8%. As seen for other sigmaviruses [44,45,49], previous work found DimmSV was vertically transmitted, with no evidence of horizontal transmission [50]—although that study was not specifically designed to detect it. Similarly, galbut virus is reported to be primarily vertically transmitted [17]. Our finding may thus represent evidence of a dual horizontal and vertical transmission mode for these viruses. For DimmSV, the small amounts of *D. immigrans* RNA contaminating our pooled RNA sequencing on the six positive Dmel females makes it possible that these apparent transmissions were a false positive. Additionally, we found no evidence of DimmSV reads in non-Dimm pooled sequencing (Fig S14) from three previous studies [8,9,48]. For galbut virus however, two lines of evidence support possible transmission and replication. First, flies were only exposed to contamination for three days, but RNA remained at death - a mean of 23.4 days later, and after a minimum of three passages onto new media. This is consistent either with horizontal transmission, or virus replication in embryos fathered by infected wild males. Second, in other work we observe galbut virus infecting five non-Dmel *Drosophila* species at 1-5% prevalence in wild populations around Edinburgh [38], consistent with transmission in nature. We also found very small numbers of galbut reads (22 galbut reads vs 6933 Dimm CO1) in pooled Dimm sequencing from another previous study (Fig S14). Therefore, for galbut virus, evidence of horizontal transmission remains equivocal.

### 4.3 Viral infection is associated with reduced lifespan and offspring production in laboratory Drosophila

Infection with galbut virus, Muthill virus, DimmNV, DmelNV or Prestney burn virus was associated with a 10-30% reduction (or more) in lifespan for *D. melanogaster* females (Fig. 3 & Table S4). The impact of galbut virus infection on lifespan and fecundity has been investigated previously [22]. However, no significant reduction in lifespan was identified, contrary to our results. The lifespan of flies varied significantly by line in this study, suggesting that this effect could interact with genotype. Our results concur with this previous study’s finding that galbut virus does not significantly reduce total lifetime offspring production. Muthill virus, DimmNV and DmelNV *were* associated with a significant decrease in lifetime offspring production in *D. melanogaster* females (Fig. 4 & Table S5). This is the first evidence that these three viruses reduce lifespan and offspring production, as no viral isolates are yet available.

The other virus associated with a decrease in lifespan and offspring production—DmelNV— *has* been isolated for study. Previous studies found that on DmelNV infection, flies initiate an immune response [51,52], show histology and gene expression consistent with damage to the gut [25], and an increased diversity of the gut microbiome [53]. However, our study additionally shows an association between DmelNV infection and reduced fecundity (95% CI 29 to 63% reduction in offspring production), and one which cannot be attributed solely to a decrease in lifespan (Fig. S7 & Table S6). We also describe a clear association between shorter lifespan (95% CI 6 to 28% reduction) and DmelNV infection. This is supported by a recent study showing DmelNV infection in some laboratory stocks can cause >40% lifespan reduction [54]. However, other reports of the effect of DmelNV on lifespan are mixed [25,26,53]. It’s possible that DmelNV pathology accelerates in later life in a way that is not detectable some mortality analyses [eg. 26]. The fitness consequences of DmelNV infection could also be subtle, vary based on genotype, or affect other phenotypes such as locomotor activity [26] or sleep [55].

### 4.4 Caveats

We show a strong association between the presence of viral RNA at time of death and a reduction in lifespan and offspring production. However, there are three possible opportunities for bias. First, because we assessed total offspring production by counting emerging adults onward transmission to partners or offspring will double-count the impact of infection on offspring production—attributing pre-adult lethality or reduced male fecundity as costs to female fecundity. We therefore selected one virus (DmelNV) to test the scale of such an effect by assaying partners and offspring for infection. This identified 16 additional infections, increasing the estimated transmission rate from 37.5% to 50.8%. This suggests that undetected transmission could have been relatively common, and that we cannot attribute the reduced offspring production of infected females solely to costs suffered by the female.

Second, if flies clear viral infections during their lives [see 56, and above] then this will downwardly bias our estimates of fitness costs, as flies that experienced an early-life cost no longer carry that virus at their death. For example, a previous study reported that levels of DmelNV RNA in the faeces of infected flies remained high over a 19 day time course, or dropped dramatically over 4 days and then stayed low [25]. To quantify the size of this effect, we reclassified as infected females who tested negative for DmelNV, but whose male partners or offspring (above) tested positive. For DmelNV, this lead to a very slight reduction in the impact of infection on lifespan (posterior mean = -4.82, vs. -7.82 in original model) and non-zero offspring production (poisson regression coefficient = -0.412, vs. -0.614), although it was not significantly different from using female assays alone (posteriors overlapped in both cases –S1, and Fig. S8 & S9). If the impact of transient early infection is small compared to lifelong (or life-limiting) infection this could explain this small effect.

Third, and potentially more seriously, if flies do regularly clear infection during their lives, then this could generate a spurious association between infection and survival. This is because flies that die early get tested earlier, potentially generating a correlation between early death from other causes and a positive virus assay. This could also affect early-life offspring production, if longevity and early-life fecundity are positively correlated. We therefore tested whether exposure *per se*, regardless of detectable virus transmission affected longevity or offspring production, reasoning that the only compelling explanation for differences in fitness-associated traits 30-40 days post-exposure must be a transfer of microorganisms (see S1 for methods and results). The three day exposure of laboratory females to wild (but not laboratory) Dmel was associated with a significant reduction in lifespan and offspring production, and exposure to wild Dimm with a significant reduction in lifespan (Fig S10 & S12, Table S7 & S8). This strongly supports the real effect of transmitted microbiota on lifespan and offspring production.

### 4.5 Power and the strength of selection

For several viruses, we saw no significant impact on fitness-related traits. For many viruses this likely reflects low power, but for some with no suggestion of a reduction in fitness, it might suggest that such viruses are commensal or even mutualistic [57–59]. However, such a conclusion is difficult to reach with the observed mean and residual variances as around 45 thousand flies per treatment group would be required to detect a 1% reduction in lifespan (S1 & Fig S13). And, given the large effective size of *Drosophila* populations, fitness costs (or benefits) of 0.1% or even 0.01% could drive host evolution.

Nevertheless, despite relatively small sample sizes, we found substantial reductions in longevity and offspring production in *Drosophila* naturally infected by several viruses. This is consistent with the scale of the mortality increase seen in larvae orally infected with DCV [60,61], with the lifespan reduction seen in some DmelNV infected Dmel stocks [27], and with the fecundity reduction seen in susceptible Dmel orally infected with DAV [4], and in *D. suzukii* orally infected with Newfield virus [18]. Given the large effective population size of *Drosophila* species, such reductions in fitness—around several or even tens of percent—could represent enormous selective pressures if translated to the wild. We lack comparable wild estimates for many other *Drosophila* viruses, but infection with Dmel sigma virus is estimated to reduce fitness by ∼20-30% in the wild [23,62]. Even more strikingly, reductions in lifespan (58%) and offspring production (70-80%) in wild *D. innubila* with Drosophila innubila Nudivirus far exceed our observed reductions in fitness [6]. This suggests that this scale of virus-driven selective pressures might frequently be present in *Drosophila* populations.

## Supporting information

Supplemental material details

Supplementary methods and results

## Competing interests

We declare we have no competing interests

## Funding

This work was funded by the UK Natural Environmental Research Council through an E3 doctoral training programme (NE/L002558/1) studentship to M.W. For the purpose of open access, the authors have applied a Creative Commons Attribution (CC BY) licence to any Author Accepted Manuscript version arising from this submission.

## Acknowledgements

We thank the City of Edinburgh Council, the Friends of the Hermitage of Braid and Keith and Sue Obbard for permission to collect flies, and we thank Jarrod Hadfield for statistical advice.

## Notes

https://doi.org/10.6084/m9.figshare.22324042.v3

### Competing Interest Statement

The authors have declared no competing interest.

