## Supplemental material details for "Naturally occurring viruses of *Drosophila* reduce offspring number and lifespan"

### Details of supplementary material

- **S1 - Supplementary methods.** OreR Wol- stock creation, Drosophila medium recipes, RT-PCR methods, Sanger sequencing and phylogeny building, analysis of public sequencing datasets, power simulations, and analyses with male, offspring and exposure species data.

##### Tables

- **Table S1 – Table of primers used to assay wild-collected flies for viruses.** Details of primer assays and conditions used to scan pools of wild-collected D. immigrans, *D. melanogaster* and obscura group species for viruses.
- **Table S2 – Table of primers used to assay female *D. melanogaster* OreR.** Details of short-product primer assays and conditions used to scan OreR Wol- females who were exposed to viruses carried by wild-collected flies. All assays used an extension time of 1 minute.
- **Table S3 – Table summarising the virus transmission results.** For viruses detected in the wild-collected flies by RT-PCR, the prevalence of these viruses was estimated in each wild species using maximum likelihood, taking into account the number of pooled Bernoulli trials, the size of the pools, and the no. of 'successes' (positives) in those pools. We quantified transmission into *D. melanogaster* using the number of individuals infected with a virus as a proportion of the number exposed, along with binomial confidence intervals.
- **Table S4 – Table containing outputs from linear mixed effects models used to analyse the effect of viral infection on lifespan in female Dmel.** Fixed effect predictors were determined to be significant if their 95% posterior credible interval did not overlap zero.
- **Table S5 – Table containing outputs from the hurdle-Poisson mixed effects model used to analyse the effect of viral infection on lifetime offspring production in female Dmel.** The probability of zero offspring was only allowed to vary with infection for eight viruses, as no zero-offspring individuals were infected with DimmSV or Corseley virus.
- **Table S6 – Table containing outputs from the hurdle-Poisson mixed effects model used to analyse the effect of viral infection on early life offspring production in female Dmel.** The probability of zero offspring was only allowed to vary with infection for five viruses, as in the early life offspring dataset, five of the viruses infected no zero-offspring individuals.
- **Table S7 - Table containing outputs from a linear model used to analyse the effect of exposure to wild/lab flies on lifespan in female Dmel.** The fixed effect

predictors (exposure species – four levels) were determined to be significant if their 95% credible interval did not overlap zero.

- **Table S8 – Table containing outputs from the hurdle-Poisson model used to analyse the effect of exposure species on lifetime offspring production in female Dmel.** This model examined the effect of being exposed to each of the wild species groups on lifetime offspring production, compared to being exposed to lab OreR Dmel. The model examines the effect of exposure on 1) non-zero offspring production, and 2) the probability of zero-offspring production.

#### Figures

- **Fig. S1 – Predicted number of *D. melanogaster* females with zero offspring under chosen hurdle model for lifetime offspring production.** The frequency histogram shows the distribution of predicted zero-offspring values when the offspring data was simulated 3000 times under the hurdle model. The yellow line indicates the actual number of zero-offspring individuals recorded in our data (25 females).
- **Fig. S2 - DimmNV maximum clade credibility tree including experimental and wild collected flies.** The tree shows the phylogenetic relationships between a ~600pb region of VP2 (RdRp) of DimmNV. Host flies were from donor and recipient infections in the experiment, and wild-collected *Drosophilidae* from the South of Scotland. Branches are coloured by host species, node circles are present if posterior support >= 0.6, and tip labels display the host species and collection country. The three highlighted (red) tip labels are DimmNV infections from this experiment, 2 donor vials (1 Dmel and 1 Dimm), and one recipient OreR Dmel.
- **Fig. S3 - Phylogenies of ImmSV sequences (regions of the L and N genes) from experimental and wild collected flies.** The trees show the phylogenetic relationship between a ~625 bp region of the L (RdRp) and a ~525bp region of the N (nucleocapsid) genes of DimmSV. Host flies were from this experiment, wild-collected *Drosophilidae* from Southern Scotland, and collections made as a part of other studies [50,63,64]. Branches are coloured by the most likely ancestral host species, node circles are present if posterior support >= 0.6, and tip labels display the host species and collection country. The highlighted (red) tip labels are DimmSV infections from this experiment, including donor vials (wild-collected flies), and experimental recipients (female Dmel OreR).

- **Fig. S4 - Figure showing a comparison between the outputs from two models of lifespan in *D. melanogaster* females.** The plot shows the lifespan of *D. melanogaster* females when clear of viruses (grey), and when infected by ten different viruses (purple). Boxplots and points indicate the raw data with median and inter-quartile range. Solid horizontal lines and points indicate the model predictions and 95% HPD credibility intervals from the fixed effects (black) and random effects (blue) models. Sample size indicates the number of individuals that tested positive for each virus. More details on model specification can be found in the methods, but the main differentiating factor is the specification of virus as a fixed or random effect.
- **Fig. S5 - Figure showing the effect of the number of viral infections on lifespan in female Dmel.** The plot shows the distribution of female lifespan observed when uninfected (grey), singly, or multiply infected with viruses (purple). Boxplots and points indicate the raw data with median and inter-quartile range. Solid horizontal lines and points indicate the model predictions and 95% HPD credibility intervals from the 'viruses as random effects' model, in which number of viral infections was a significant predictor of lifespan. Sample size indicates the number of *D. melanogaster* females in each category.
- **Fig. S6 - No. of offspring produced by *D. melanogaster* females, over time, by infection status.** Panels show the offspring produced over time by infected (purple) and uninfected (grey line) laboratory *D. melanogaster* females. The number of infected flies is shown for each virus (n) and the points indicate the mean offspring per day (bars give ±1 standard deviation around the mean). Note that the same 393 uninfected flies are depicted in each graph for comparison. Only those viruses with at least 10 infections are included.
- **Fig. S7 – Figure showing a comparison between posterior density distributions for viral fixed effects included in the model of lifetime offspring production, and early life offspring production.** Filled curves show the posterior density of estimates displayed in purple (zero-truncated Poisson) and blue (probability of zero offspring) for the lifetime offspring production hurdle model, and in yellow for the model of early life offspring production, with solid horizontal lines and points to indicate the model prediction and 95% HPD credibility intervals.
- **Fig. S8 – Figure showing posterior density distributions for viral fixed effects included in the model of lifespan variation, with extra data from males and offspring for DmelNV.** Rows show the offspring production of D. melanogaster females that either were uninfected (top row), or infected by one of ten viruses (subsequent rows). Filled curves show the posterior density of estimates displayed in

purple/grey for the original lifespan model, and the distribution of the effect of DmelNV infection when offspring and male infection data is included is shown in yellow, with solid horizontal lines and points to indicate the model prediction and 95% HPD credibility intervals. Boxplots and points indicate the raw data with median and inter-quartile range. Sample size indicates the number of Dmel individuals in each category, with numbers in brackets indicating sample size with the extra data from males and offspring included for DmelNV.

- **Fig. S9 – The impact of viral infection on Dmel offspring production, with extra data from males and offspring for DmelNV.** Rows show the offspring production of D. melanogaster females that either were uninfected (top row), or infected by one of ten viruses (subsequent rows). The left side of the plot shows the effect of exposure on non-zero offspring production. Boxplots and points indicate the raw data with median and inter-quartile range. Filled curves show the posterior density of estimates, with solid horizontal lines and points to indicate the model prediction and 95% HPD credibility intervals. Sample size indicates the number of individuals that were exposed to each treatment. The right side of the plot shows the predicted probability of zero offspring being produced, as horizontal bars. Dashed line boxes indicate the 95% HPD credibility intervals on these estimates. The original model distributions are displayed in purple/grey, and the distribution of the effect of DmelNV infection when offspring and male infection data is included is shown in yellow.
- **Fig. S10 - Posterior estimates of lifespan by exposure species.** Rows show the lifespan of D. melanogaster females that either were exposed to OreR Dmel from the lab (the control - top row), or three species groups of wild flies (subsequent rows). Boxplots and points indicate the raw data with median and inter-quartile range. Filled curves show the posterior density of estimates, with solid horizontal lines and points to indicate the model prediction and 95% HPD credibility intervals. Sample size indicates the number of individuals that were exposed to each species group/control.
- **Fig. S11 - Predicted number of *D. melanogaster* females with zero offspring under a hurdle model including exposure species, for lifetime offspring production.** The frequency histogram shows the distribution of predicted zero-offspring values when the offspring data was simulated 3000 times under the hurdle model. The yellow line indicates the actual number of zero-offspring individuals recorded in our data (25 females).
- **Fig. S12 - The impact of exposure to wild/lab flies on lifetime offspring production in female *D. melanogaster* (OreR).** Rows show the offspring production of *D. melanogaster* females that either were exposed to Dmel OreR from the lab

(control - top row), or one of three species groups of wild flies (subsequent rows). The left side of the plot shows the effect of exposure on non-zero offspring production. Boxplots and points indicate the raw data with median and inter-quartile range. Filled curves show the posterior density of estimates, with solid horizontal lines and points to indicate the model prediction and 95% HPD credibility intervals. Sample size indicates the number of individuals that were exposed to each treatment. The right side of the plot shows the predicted probability of zero offspring being produced, as horizontal bars. Dashed line boxes indicate the 95% HPD credibility intervals on these estimates.

- **Fig. S13 – Simulations of power to detect a viral-induced reduction in lifespan, with different sample sizes.** Plots show the sample size needed to detect a A) 20%, B) 10%, C) 5%, and D) 1% reduction in lifespan due to infection by a virus. Each data point represents the mean power (over 1000 simulations) to detect the reduction in lifespan, and how this changes as sample size is increased within each experimental group. The line indicates smoothed conditional means for these points. The simulations were created using the mean and variance in lifespan measured from this experiment.
- **Fig. S14 – Heat map of the number of viral genome copies or host CO1 transcripts in pooled *Drosophila* sequencing from three previous studies.** The heatmap shows the number of reads from each sequencing pool which map to either a version of the galbut virus genome, DimmNV or DimmSV, or any Drosophila CO1 gene. Read numbers are normalised by target sequence length. The raw numbers – which can be found on our Figshare repository – were used to assess whether these viruses can infect non-Dmel, or non-Dimm species respectively. Data was downloaded from https://ftp.sra.ebi.ac.uk/, more details in S1.
