## Supplementary methods and results for "Naturally occurring viruses of *Drosophila* reduce offspring number and lifespan"

### S1. Supplementary methods and results

#### OreR Wolbachia$^-$ Stocks

To provide stocks of laboratory Dmel, we set up bottles of age matched OregonR *Wolbachia* negative flies (OreR Wol-) using egg 'squirts' (eggs collected from agar and suspended in pbs) to limit the faecal-oral spread of viral infection between adult *Drosophila* and larvae, and to control density. Samples of 10 OreR males were frozen from each of the recipient stock bottles to test for contaminating viral infections using later PCR assays.

#### RNA Extractions, and RT-PCR methods

RNA was extracted from the wild donor flies and single OreR Wol- flies using a manual Phenol-Chloroform protocol. Briefly, the flies were macerated in Tri Reagent solution (ThermoFisher Scientific) (100µL for single flies, 200uL for pools of 10) and RNA extracted using the manufacturer's instructions. cDNA from each sample was synthesised using random hexamer primers (ThermoFisher Scientific; 4µL 10 mM random primers per 1µL total RNA) which were then incubated with the RNA (70ºC, 5 minutes). RNase free $H_2O$, 10 mM mixed dNTPs (ThermoFisher Scientific), M-MVL reverse transcriptase (200 units/µl, Promega) and 5x M-MVL reaction buffer (Promega) were then added to the 5µL RNA-random primer mix and incubated at 37ºC for 60 minutes.

To reveal the viruses to which the female OreR had been exposed during co-housing, we tested each of the wild collected donor groups of *Drosophila* (Dimm, Dobs and Dmel) and the control OreR males for the presence of a total of 59 different viruses by RT-PCR (primers listed in table S1). Additional PCR primers with relatively short product lengths can be found in table S2. All PCRs were performed with a master mix of the following volumes per 1µl of cDNA template - 1µl 10xNH4 buffer, 0.25µl 50mM MgCl2, 0.05µl 5U/µl BIOtaq DNA polymerase (all Bioline reagents ltd.), 0.3µl 10mM mixed dNTPs (Life Technologies), 7.5µl

RNase free H2O, and 0.5µl of both 10µM forward and reverse primers (Sigma Aldrich). All PCRs were run on a thermocycling regime of 94ºC for 5 minutes, then 5-10 x (94ºC for 15s, * ºC for 30s, 72 ºC for 1 minute) dropping $1^0$C every cycle, then 25 x (94ºC for 15s, * ºC for 30s, 72 ºC for 1 minute), then 72 ºC for 5 minutes, with the * ºC adjusted for virus assays. The Tm for each primer pair can be found in table S1 & S2. Presence or absence of a virus was determined using gel electrophoresis, stained using GelRed 10,000x (Biotium), on a 1 or 2% agarose gel (dependant on the size of the PCR products) at 90V.

#### Sanger sequencing of DimmNV and DimmSV

The products of successful RT-PCR reactions targeting DimmSV and DimmNV infections were treated with Exo-SAP-IT ® Express Reagent (Applied Biosystems) according to the manufacturer's instructions. 4µL (recipients) or 1µL (donors) of the resulting product were then sequenced by adding 1µL BigDye reagent from BigDye ® Terminator v3.1 Cycle Sequencing Kit (Applied Biosystems), 0.7 µL 3.2mM primers and RNase free $H_2O$ (volume to make the total reaction 10 µL) and incubating for 1 minute at 96ºC, then 25x (10s at 96ºC, 5s at 50 ºC, 4 minutes at 60 ºC). Capillary electrophoresis was performed by Edinburgh Genomics (Edinburgh). The resulting sequences were quality trimmed at ends, and pools with multiple flies were examined for the presence of heterozygous sites. A BLAST search then identified the closest virus sequence in NCBI databases [1]. The viral sequences, additional sequences from other studies, and sequences from wild collected *Drosophila* collected in Scotland [2] were then aligned using a codon model in PRANK multiple sequence aligner [3]. These alignments were used to construct phylogenies of the virus sequences, whilst inferring the most likely host species on ancestral nodes, in BEAST v1.10.4 [4]. We used the SDR06 substitution model [5], and default priors, other than a lognormal prior on the strict clock rate with a mean and initial value of $5x10^{-4}$, and standard deviation of $2x10^{-4}$ (a reasonable rate for RNA viruses). Estimates of population growth rate in preliminary models showed that a constant population size tree coalescent was most appropriate. For the DimmSV trees, we allowed ancestrally reconstructed host species transmission rates to vary independently, such that the rate of transmission from Dimm to Dmel was allowed to differ from the reverse direction of transmission. Two Markov chain Monte Carlo (MCMC) chains with a length $5x10^7$ were run, sampling 10,000 trees on each chain. We examined trace files for sufficient effective sample size and chain convergence before combining runs and generating a maximum clade credibility (MCC) tree in TreeAnnotator v1.10.4 [6], setting the burn-in to 10% of the total MCMC chain length. The MCC trees were viewed in FigTree v1.4.4 (http://tree.bio.ed.ac.uk/) and then annotated in R v.4.0.3 [7] using the ggtree package [8].

#### Mapping of public RNA sequencing data to investigate Drosophila virus host range and transmission

We analysed three publically available short read illumina sequencing datasets [9–11] to investigate unexpected presence of DimmNV, DimmSV and galbut virus infection. Data was downloaded from https://ftp.sra.ebi.ac.uk/ and can be found in the sequence read archive under Bioproject numbers PRJNA728554, PRJNA402011 and PRJNA312496. We competitively mapped the raw paired-end reads from each of the datasets back to a database containing all currently published (and some unpublished) *Drosophila* viruses from https://obbard.bio.ed.ac.uk/data.html, and *Drosophila* cytochrome oxidase 1 sequences using bowtie2 v2.4.5 [12]. We used the –very-sensitive option to reduce cross mapping between closely related viruses, and similar CO1 sequences, and quality filtered mapped reads at –q 10 using samtools v1.14 [13]. To get a measure of the abundance of viral genome copies or host CO1 transcripts, we counted reads mapping to each virus or host gene and normalised these counts by contig length. We plotted these results in a heatmap produced using the R package pheatmap [14].

#### Simulations of the sample size needed to detect lifespan reduction

We had insufficient power to detect fitness costs for many of the transmitted viruses in this experiment. To assess the sample size that would be needed to detect different effect sizes on lifespan, we ran power simulations using the R package simglm [15].  In these simulations we varied sample size to a maximum of 100,000 flies per treatment, and the mean reduction in lifespan caused by a virus between 1% and 20%. These simulations assume a mean (41.9) and variance (266.7) in lifespan as measured from this experiment, and a balanced experimental design with only one viral infection.

#### The impact of exposure species on Dmel lifespan and offspring production

As an initial assessment of the effect of exposing Dmel OreR females to wild flies of three species groups, we ran linear models in the Bayesian package MCMCglmm [16] in R v.4.0.3 [7]. We examined the effect of species of exposure on lifespan, and lifetime offspring production (relative to being exposed to OreR males from the lab as the control). Exposure species was fitted as a fixed effect with four levels in both models. Lifespan was modelled using a Gaussian distribution. The sample size of the Markov chain was 10,000, and all estimates had an effective sample size of >9,500. To account for zero-inflation in the data, lifetime offspring production was modelled using a hurdle-Poisson distribution (see main text). For the offspring model, we used simulations based on our posterior estimates to confirm that our estimated parameters were consistent with the observed number of zeros (Fig. S11).The sample size of the Markov chain was 10,000, and all estimates had an effective sample size of >2,500.

Being exposed to wild Dmel, or wild Dimm, was associated with a significant reduction in lifespan (compared to the control). Exposure to Dmel resulted in a mean lifespan reduction of 6.25 days (HPD -10.94 to -1.80), while exposure to Dimm reduced lifespan by 4.81 days (HPD -9.09 to -0.60) (Table S7, Fig. S10). Being exposed to wild Dmel was also associated with reduced lifetime offspring production (Poisson regression coefficient -0.34, HPD -0.605 to -0.057; pMCMC = 0.017) (Table S8, and Fig. S12).

#### The impact of viral infection on Dmel lifespan and offspring production – analysing infections in males and offspring

It is possible that flies may clear infections prior to death, causing some infections to be missed, such as seen in DCV [17]. To test the impact of this on our inferences, we selected one common and highly transmissible virus (DmelNV) and additionally tested males and offspring that may have received the virus from an infected female who later cleared the infection. The males housed with the virus-exposed females were frozen at -80C on death of the female; i.e. before their own death. We also froze offspring produced in the first 2 days of the experiment. Therefore, by taking not only female infections, but also male and offspring infections into account, we could detect DmelNV infections that would otherwise be missed through infection clearance in the focal female or RNA degradation subsequent to death. We then included the male and offspring infections in the data by assigning a female Dmel individual as infected with DmelNV if either she, or her offspring or accompanying males were infected (16 additional infections). We analysed this data using the '*viruses as fixed effects'* model for lifespan and offspring production; we used the same hurdle-Poisson model as in the main analysis (see main text). For the model of lifespan, all posterior estimates had an effective sample size >5,000, and for lifetime offspring production, >2,000.

We found that when data on infections in accompanying males and offspring was included in the analysis of lifespan, it did not change the overall results (Fig. S8). The mean effect size of DmelNV was reduced (posterior mean = -4.82, vs. posterior mean = -7.82 in original model), though the posteriors overlapped (HPD -9.42 to -0.31, HPD -12.93 to -2.77 in original model). The mean effect of DmelNV infection on non-zero offspring production was also reduced (Poisson regression coefficient = -0.412, vs. -0.614 in original model), but again, the posteriors overlapped (HPD -0.7 to -0.13, HPD -0.94 to -0.3 in original model). As in the original model, there was no effect of DmelNV infection on the probability of zero offspring production (Fig. S9).

#### Lewis medium recipe

| **Ingredients** | **Quantity** (e.g. for 300 vials/100 bottles) |
|---|---|
| 3x dd. $H_2O$ (l) | 3.75 |
| Agar (g) | 23 |
| Brown sugar (g) | 351 |
| Maize (g) | 255 |
| Yeast (g) | 70 |
| Nipagen (ml) | 56 |

- Mix all ingredients apart from Nipagen in a pan and put on the hob, stir until boiling (10 mins), and then turn down heat once boiled and continue stirring for 5-10min/until reached desired consistency
- Once mixture cooled to 70°C, add Nipagen, and check pH of mixture using litmus strips. Needs to be pH7, can adjust by adding 5ml of sodium hydroxide at a time until correct pH
- Dispense into vials or bottles and leave to cool overnight

#### Agar/sugar medium recipe

| **Ingredients** | **Quantity** (e.g. for 300 vials/100 bottles) |
|---|---|
| 3x dd. $H_2O$ (l) | 1 |
| Agar (g) | 20 |
| Brown sugar (g) | 84 |
| Tegosept/Nipagin (ml) | 7 |

- Mix agar, brown sugar and $H_2O$ in a pan. Place on heat and bring the mixture to boil while stirring continuously.
- Take off heat and allow mixture to cool to ~70 °C and then stir in tegosept/nipagin
- Dispense agar into vials before it sets.
